# IS*1*-related large-scale deletion of chromosomal regions harbouring oxygen-insensitive nitroreductase gene *nfsB* causes nitrofurantoin heteroresistance in *Escherichia coli*

**DOI:** 10.1101/2023.04.03.535367

**Authors:** Yu Wan, Akshay Sabnis, Zaynab Mumin, Isabelle Potterill, Elita Jauneikaite, Colin S. Brown, Matthew J. Ellington, Andrew Edwards, Shiranee Sriskandan

## Abstract

Nitrofurantoin is a broad-spectrum first-line antimicrobial used for managing uncomplicated urinary tract infection. Loss-of-function mutations in chromosomal genes *nfsA, nfsB*, and *ribE* of *Escherichia coli* are known to reduce nitrofurantoin susceptibility. Here, we report monoclonal nitrofurantoin heteroresistance in *E. coli* and a novel genetic mechanism associated with this phenomenon.

Subpopulations with reduced nitrofurantoin susceptibility in cultures of two *E. coli* blood strains were identified using population analysis profiling. Four colonies of each strain growing on agar with 0.5×MIC nitrofurantoin were sub-cultured in broth with 0.5×MIC nitrofurantoin (n=2) or without nitrofurantoin (n=2). Moreover, one colony of each strain growing without nitrofurantoin exposure was selected as a reference for genomic comparison. Whole-genome sequencing of all isolates were conducted on Illumina and Nanopore MinION systems.

Both strains had a nitrofurantoin MICs of 64 mg/L. The proportion of cells grown at 0.5×MIC was two and 99 per million, respectively, which is distinct to that of a homogeneously susceptible or resistant isolate. All isolates grown at 0.5×MIC had 11–66 kbp deletions in chromosomal regions harbouring *nfsB*, and all these deletions were immediately adjacent to IS*1*-family insertion sequences.

Although this study is limited to *E. coli* and nitrofurantoin, our findings suggest IS*1*-associated genetic deletion represents a hitherto unrecognised mechanism of heteroresistance that could compromise infection management and impact conventional antimicrobial susceptibility testing.

**Impact statement:** Nitrofurantoin is widely used for treating and preventing urinary tract infection. Prevalence of nitrofurantoin resistance generally is low in *E. coli*. Here, we report nitrofurantoin heteroresistance in two *E. coli* blood strains and attribute this phenotype to IS*1*-associated deletion of chromosomal regions harbouring oxygen-insensitive nitroreductase gene *nfsB*. Our discoveries demonstrate a novel genetic mechanism of heteroresistance and suggest detecting nitrofurantoin heteroresistance in *E. coli* urinary isolates for improving prescribing.

**Data summary:** Whole-genome sequencing reads and genome assemblies generated in this study have been deposited under BioProject PRJEB58678 in the European Nucleotide Archive (ENA). Accession numbers are listed in Supplementary Table 1. Previously generated Illumina whole-genome sequencing reads of parental isolates EC0026B and EC0880B are available under ENA accessions ERR3142418 and ERR3142524, respectively.

## Introduction

Nitrofurantoin is a widely used first-line antimicrobial for urinary tract infection (UTI) treatment and prophylaxis [1]. Reduced nitrofurantoin susceptibility in *Escherichia coli* has been associated with deleterious mutations in chromosomal genes *nfsA, nfsB*, and *ribE* [2], which encode key components of the oxygen-insensitive nitroreductase system, and with acquired multidrug efflux pump OqxAB [3]. Despite low prevalence of nitrofurantoin resistance in *E. coli* clinical isolates (<7% in Europe) [4, 5], concerns over increased prevalence of nitrofurantoin resistance are growing in England, where the consumption of nitrofurantoin had increased by 41% from 2017 to 2021 [6].

We previously identified three *E. coli* clinical strains with apparent signs of heteroresistance to nitrofurantoin [7], a phenomenon that may easily be overlooked in clinical antimicrobial susceptibility testing. We therefore set out to test nitrofurantoin heteroresistance and explore mechanisms of this phenotype for two strains that showed similar growth patterns.

## Methods

### *E. coli* strains

Two blood strains EC0026B (SAMEA104039660) and EC0880B (SAMEA104040147) [8] were tested to confirm nitrofurantoin heteroresistance. A nitrofurantoin-resistant strain IN09 [7] and nitrofurantoin-susceptible strain ATCC 25922 [9] were included as positive and quality controls, respectively, for population analysis profiling (PAP). Illumina whole-genome sequencing (WGS) reads of parental isolates EC0026B and EC0880B have been generated previously [8] and are available under accessions ERR3142418 and ERR3142524, respectively, in the ENA. We have shown that gene *nfsA* in each of EC0026B and EC0880B carries a loss-of-function mutation [7].

### Population analysis profiling

PAP was conducted on three separate occasions for each of the four strains (Figure 1). In brief, nitrofurantoin (N7878, Sigma-Aldrich, Germany) was dissolved in dimethylformamide to produce a stock solution, which was then infused into cation-adjusted Mueller-Hinton (MH2; 90922, Sigma-Aldrich) agar to form a two-fold gradient of nitrofurantoin concentrations (4– 256 mg/L). One colony of each strain was inoculated into nitrofurantoin-free MH2 broth and grown aerobically overnight at 37 °C with shaking (180 RPM). Seven serial 10-fold dilutions of each broth culture were created with phosphate buffered saline (P4417, Sigma-Aldrich). Ten-μL aliquots of the original broth culture and serial dilutions were spread in octants of MH2 agar plates following the nitrofurantoin gradient and incubated aerobically overnight at 35 °C. For each agar plate, colony-forming units (CFUs) were counted in the octant where colonies were well separated, and thereby nitrofurantoin MIC of each strain was determined. A maximum non-inhibitory nitrofurantoin concentration of each strain was defined as the greatest concentration in which ≥70% cells grew when compared to the growth without nitrofurantoin. The European Committee on Antimicrobial Susceptibility Testing (EUCAST) clinical breakpoint (v13.0) was used for interpreting nitrofurantoin susceptibility of strains (susceptible: ≤64 mg/L; resistant: >64 mg/L) [10].

**Figure 1.**
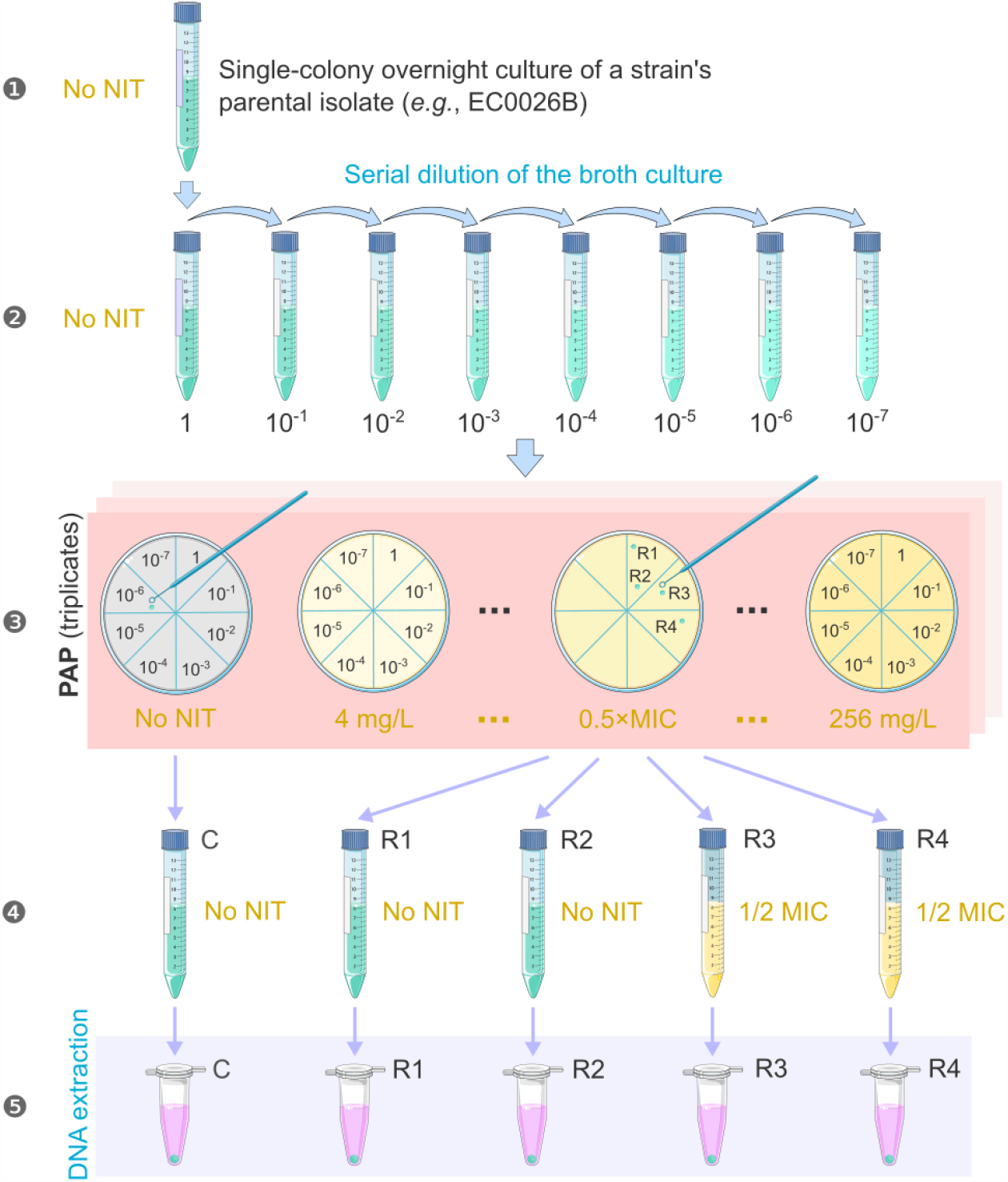
Workflow of population analysis profiling for each *E. coli* strain and preparation of isolates for DNA extraction. Isolate C represents a comparator controlled for the number of passages. Isolates R1–R4, which showed reduced nitrofurantoin susceptibility when compared to the majority of cells in the same broth culture, were randomly chosen at Step 3. Nitrofurantoin (NIT) concentrations in media are noted in gold. Icons were downloaded from Bioicons (bioicons.com) under CC-BY 3.0 Licence.

### Deriving isolates from strains EC0026B and EC0880B

Following PAP, four colonies with reduced nitrofurantoin susceptibility were randomly chosen (denoted by subscript ‘R’, *e*.*g*., EC0026B_R1_) from an agar plate containing 0.5×MIC nitrofurantoin as the exposure group for each strain. Then these isolates were evenly divided into two subgroups, inoculated into MH2 broth with or without nitrofurantoin (Figure 1), and aerobically incubated overnight at 35 °C. A colony of each strain was randomly selected from a nitrofurantoin-free agar plate as a comparator (denoted by subscript ‘C’, *e*.*g*., EC0026B_C_) to the exposure group and was grown in nitrofurantoin-free MH2 broth before DNA extraction.

### DNA extraction and whole-genome sequencing

For each isolate derived from a parental isolate (EC0026B or EC0880B), genomic DNA was extracted from the broth culture using Proteinase K Solution, RNase A Solution, and Cell Lysis Solution (QIAGEN, Germany), and purified using GeneJET Genomic DNA Purification Kit (ThermoFisher Scientific, UK). The mean concentration of DNA in each extract was estimated from three reads obtained with Qubit dsDNA BR Assay Kit (ThermoFisher Scientific).

Extracted DNA of each isolate was aliquoted for WGS. Short-read sequencing (101 bp, paired-end) was conducted on an Illumina HiSeq 2500 system (Illumina, USA), and long-read sequencing was conducted on a MinION R9.4.1 flow cell (Oxford Nanopore Technologies, UK) for isolates having adequate yields of DNA. For MinION sequencing, DNA libraries were prepared using Rapid Barcoding Kit SQK-RBK004 (Oxford Nanopore Technologies), and base-calling, demultiplexing, and barcode trimming were conducted with Guppy v5.0.16 (community.nanoporetech.com/downloads) and its built-in high-accuracy model. For quality control, Illumina reads (101 bp, paired-end) were trimmed for a minimal base quality of Phred Q20 (10-bp sliding window) and filtered for a minimal length of 50 bp using Trimmomatic [11]; MinION reads were filtered for a minimal per-read average quality of Q10 and minimal length of 1 kbp using NanoFilt v2.8.0 [12].

### Genome assembly and annotation

For isolates having MinION and Illumina reads, MinION reads were assembled *de novo* using Raven v1.7.0 [13] and polished with the same reads for four times by Raven and for an additional round by Medaka v1.7 (github.com/nanoporetech/medaka); for isolates only having Illumina reads, genomes were assembled using Unicycler v0.5.0 [14]. All assemblies were then polished with Illumina reads using Polypolish v0.5.0, POLCA v4.0.9, and again Polypolish [15–17]. Complete chromosome and plasmid assemblies were rotated to start from *dnaA* and *rep*, respectively. Scripts used for aforementioned steps are available at github.com/wanyuac/Assembly_toolkit. Genome annotation was conducted with Prokka v1.14.6 [18]. Insertion sequences were inferred from annotations and searched against the ISFinder database [19] for confirmation.

### Variant identification

For each strain, the complete chromosome sequence of the comparator isolate was used as a reference to identify chromosomal genetic variation in isolates R1–R4. Since chromosomes of isolates having MinION reads were fully assembled, strain-specific alignments of these chromosomes against references were performed using Minimap2 v2.24 [20]. Chromosomal mutations and structural variants were identified from alignments using paftools.js (Minimap2). For isolates only having Illumina reads, chromosomal mutations were identified from strain-specific read mapping using Minimap2, Samtools v1.16.1, and BCFtools v1.16 (DP ≥10 & QUAL ≥20 & MQ ≥30) [21], and structural variants were determined with breseq v0.37.1 [22]. For each strain, mutations were filtered to exclude those in repetitive chromosomal regions (≥90% nucleotide identity, determined in the reference with MUMmer v4.0.0rc1) [23] for accuracy and then mapped against annotations of the reference sequence with SnpEff v4.3.1t to estimate functional impacts [24]. Mutations and structural variants were inspected in sequence alignments using Artemis v18.2.0 [25], and genetic structures were illustrated using BRIG v0.95 [26] and R package gggenes (wilkox.org/gggenes).

## Results

### Phenotypic confirmation of nitrofurantoin heteroresistance

Population analysis profiling (PAP) confirmed monoclonal nitrofurantoin heteroresistance in two strains EC0026B (Achtman ST484) and EC0880B (Achtman ST58). Each heteroresistant strain had a nitrofurantoin MIC of 64 mg/L, which was 8–16 times its maximum non-inhibitory concentration (EC0026B: 4 mg/L; EC0880B: 8 mg/L; Figure 2). The average proportion of cells growing at 0.5×MIC was 9.85×10^−5^ and 2.14×10^−6^ for EC0026B and EC0880B, respectively. In comparison, the MIC of control strain ATCC 25922 (16 mg/L) was four times its maximum non-inhibitory concentration, and positive control IN09 (MIC = 128 mg/L) showed a drastic transition from full growth to complete inhibition when the nitrofurantoin concentration doubled from 64 mg/L.

**Figure 2.**
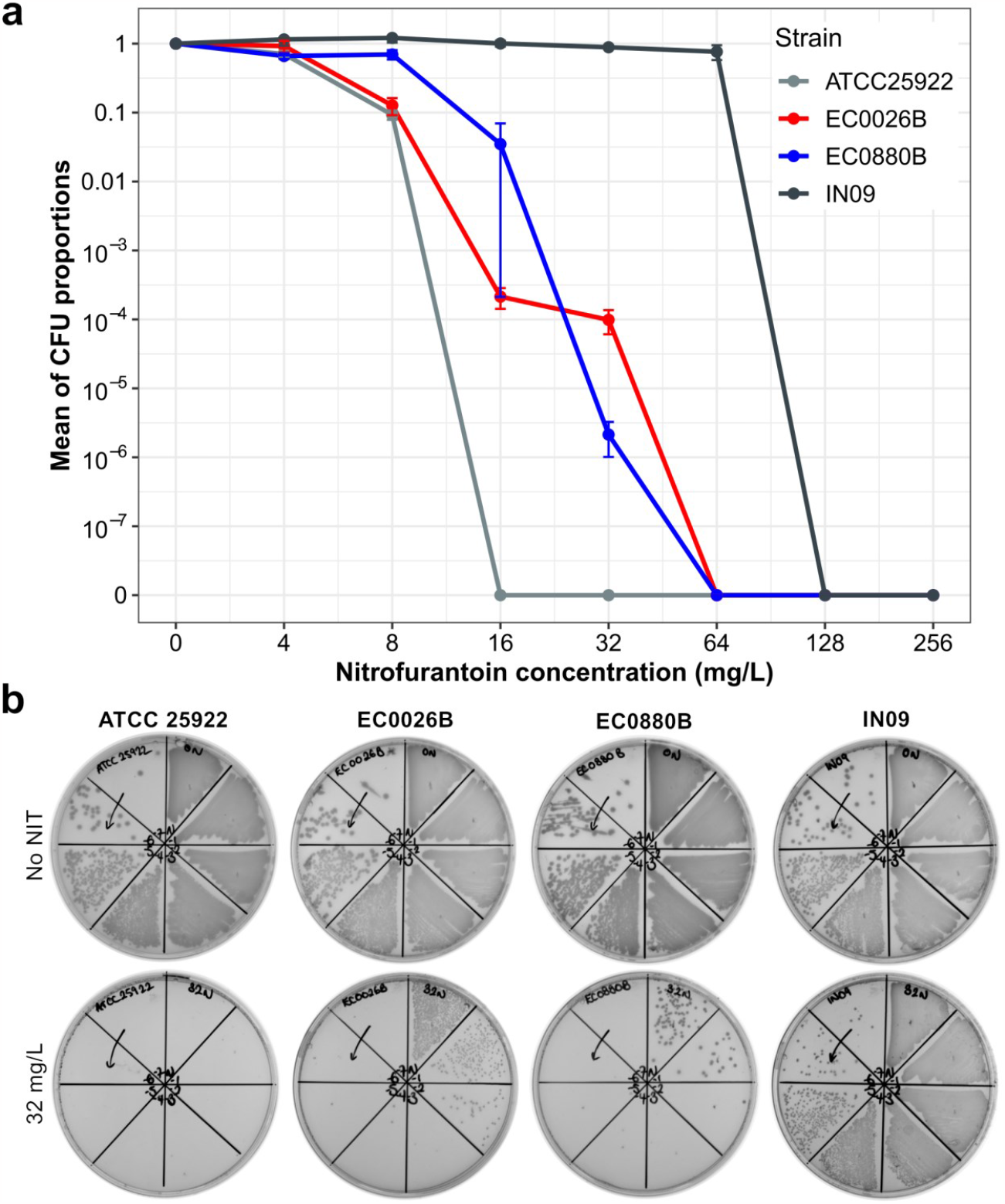
(**a**) Curves of population analysis profiling for test and control *E. coli* strains. Test strains EC0026B and EC0880B demonstrate nitrofurantoin (NIT) heteroresistance. Control strain IN09 is known to be NIT-resistant, and ATCC 25922 is susceptible. For each strain, the proportion of colony-forming units (CFUs) growing at each NIT concentration (4–256 mg/L) was calculated by dividing the CFU with the inoculum CFU (counted from the NIT-free agar plate). Based on biological triplicates, the mean proportion of CFUs ± the standard error of the mean is shown. CFUs below the detection limit are denoted by ‘0’ on the Y axis. (**b**) Growth of test and control strains on cation-adjusted Mueller-Hinton agar plates with or without NIT on the same occasion. Test strains EC0026B and EC0880B showed subpopulations growing at 32 mg/L of NIT.

### *E. coli* isolates and genome assemblies

Five isolates (R1–R4, and C) were derived from each of strains EC0026B and EC0880B (Figure 1), providing 10 isolates altogether for WGS. Complete genomes were assembled for nine of these isolates (Supplementary Table 2).

### Genetic variation in isolates derived from strain EC0026B

Isolates EC0026B_R1–R4_ showed different deletions of 11–20 kbp chromosomal regions harbouring *nfsB* (Figure 3a), and each deletion immediately followed the right imperfect inverted repeat (IRR) of an IS*1*-family insertion sequence IS*1A*, which interrupted insertion sequence IS*150* (Figure 3b). The other end of the deletion seemed variable and occurred in coding sequences. For instance, the deletion in EC0026B_R4_ chromosome ended at 10 bp upstream of the left imperfect inverted repeat (IRL) of another IS*1A*, which interrupted gene *vgrG*. These two IS*1A* elements and the relevant reference sequence in the ISFinder database mutually differed by two nucleotide substitutions in the second open reading frame (*insB*) of the IS*1A* transposase gene [27].

**Figure 3.**
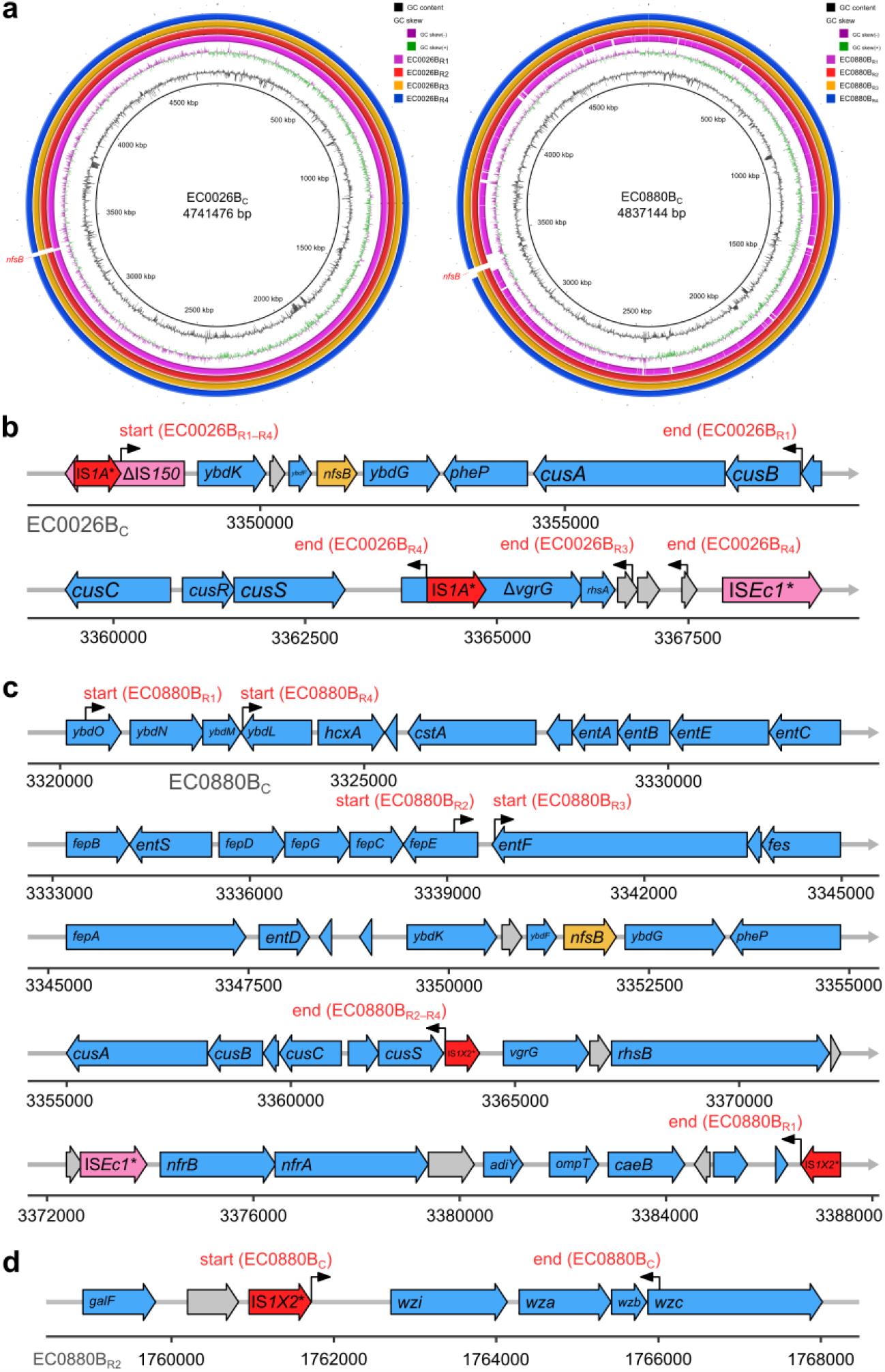
Genetic structures of deleted regions. (**a**) BRIG plots comparing genome assemblies against chromosomal sequences of isolates EC0026B_C_ and EC0880B_C_. Sequences were aligned using megaBLAST, and matches were filtered for a minimum nucleotide identity of 90% and a minimum query coverage of 0.6%. (**b–c**) Genetic structures of *nfsB*-carrying regions deleted from the chromosomes of EC0026B_C_ and EC0880B_C_, respectively, as indicated in (a). Start and end positions of each deletion are indicated by black arrows and red labels. Asterisks indicate variants of insertion sequences against reference sequences from the ISFinder database, and grey wide arrows indicate coding sequences of hypothetical proteins. (**d**) Genetic structure of the capsule-encoding region in the chromosome of EC0880B_R2_ and the partial deletion of this region in EC0880B_C_.

Twenty-four copies of IS*1A* were found in chromosomes of EC0026B_C_, EC0026B_R1_, and EC0026B_R4_, and 23 copies were found in EC0026B_R2_ and EC0026B_R3_, which is consistent with deletions illustrated in Figure 3b. These IS*1A* elements demonstrated 99–100% nucleotide identities and 100% coverage to the reference IS*1A* sequence. The explicit copy number of IS*1A* could not be determined in the Unicycler short-read-only assembly of parental isolates EC0026B. Nevertheless, in the assembly graph, the mean and median depths of assembled segments from which the complete IS*1A* was recovered suggest 20–22 copies of this element with >98% nucleotide identities to the reference sequence.

As for other chromosomal genetic variation, 14 nucleotide substitutions and two indels were found in EC0026B_R1_, whereas no point mutation was found in EC0026B_R2_, EC0026B_R3_, or EC0026B_R4_ (Table 1). No genetic variation was detected between EC0026B_C_ and EC0026B.

**Table 1.** Genetic variation in isolates EC0026B_R1–R4_ when compared to the chromosome of 26B.C. aa, amino acids.

| Isolate | Position | DNA change | Gene | Protein change |
| --- | --- | --- | --- | --- |
| EC0026B<sub>R1</sub> | 449628 | T>C | *chiA* | S779P |
|  | 461138 | A>G | *gspC* | None (synonymous) |
|  | 603693–603694 | Insertion: 78 bp | *deaD* | G585(26 aa)R586 |
|  | 1417135 | C>T | - | None (intergenic) |
|  | 1705072 | A>G | *dld* | None (synonymous) |
|  | 1962017 | T>C | *uvrY* | L176P |
|  | 1977043 | Deletion: T | *uspC* | Q18fs |
|  | 2128454 | T>G | *astD* | None (synonymous) |
|  | 2355150 | C>T | Hypothetical | V106I |
|  | 2895745 | G>A | *yccM* | A47T |
|  | 3158149 | T>C | *ybhI* | None (synonymous) |
|  | 3206276 | T>C | *gltA* | V144A |
|  | 3207019 | T>C | *gltA* | W392R |
|  | 3347713–3358894 | Deletion (11 kbp) | Multiple | Loss of protein products |
|  | 3638980 | A>G | Hypothetical | T69A |
|  | 3753165 | G>T | *aacA* | None (synonymous) |
|  | 3784606 | C>T | Hypothetical | S348N |
| EC0026B<sub>R2</sub> | 3347713–3367484 | Deletion (20 kbp) | Multiple | Loss of protein products |
| EC0026B<sub>R3</sub> | 3347713–3366768 | Deletion (19 kbp) | Multiple | Loss of protein products |
| EC0026B<sub>R4</sub> | 3347713–3364083 | Deletion (16 kbp) | Multiple | Loss of protein products |

### Genetic variation in isolates derived from strain EC0880B

Isolates EC0880B_R1–R4_ showed different large deletions (24–66 kbp) of chromosomal regions harbouring *nfsB* (Table 2). All these deletions ended immediately upstream of two identical copies of an IS*1*-family insertion sequence IS*1X2* (Figure 3a, c). However, the start site of each deletion was variable. This IS*1X2* element showed 98% nucleotide identity and 100% coverage to its reference sequence in the ISFinder database.

**Table 2.** Genetic variation in isolates EC0880B_R1–R4_ when compared to the chromosome of EC0880B_C_. Of note, the 4.3-kbp ‘insertion’ resulted from the deletion of this region in the chromosome of EC0880B_C_.

| Isolate | Position | DNA change | Gene | Protein change |
| --- | --- | --- | --- | --- |
| EC0880B<sub>R1</sub> | 1761728–1761729 | ‘Insertion’ (4.3 kbp) | *wzi, a, b, c* | No change |
|  | 3320419–3386613 | Deletion (66 kbp) | Multiple | Loss of protein products |
| EC0880B<sub>R2</sub> | 1736645 | A>T | *mdtC* | M284L |
|  | 1761728–1761729 | ‘Insertion’ (4.3 kbp) | *wzi, a, b, c* | No change |
|  | 3317674 | G>T | *ahpF* | Q147H |
|  | 3339108–3363443 | Deletion (24 kbp) | Multiple | Loss of protein products |
|  | 4117518 | A>C | *uxuB* | N187H |
| EC0880B<sub>R3</sub> | 1761728–1761729 | ‘Insertion’ (4.3 kbp) | *wzi, a, b, c* | No change |
|  | 3339723–3363443 | Deletion (24 kbp) | Multiple | Loss of protein products |
| EC0880B<sub>R4</sub> | 1761728–1761729 | ‘Insertion’ (4.3 kbp) | *wzi, a, b, c* | No change |
|  | 3323004–3363443 | Deletion (40 kbp) | Multiple | Loss of protein products |

Unexpectedly, an additional chromosomal 4289-bp deletion was identified in EC0880B_C_ when compared with that of parental isolate EC0880B despite absence of any point mutations. This deleted region was present in EC0880B_R1–R4_ (Table 2) and immediately followed the IRR of the same IS*1X2* as those related to the deletion of *nfsB* (Figure 3d). This deletion caused a loss of three genes (*wzi, wza*, and *wzb*) and a 142-bp truncation of the 5’ end of *wzc* in the conserved capsule locus *wzi-wza-wzb-wzc* [28].

Twelve copies of IS*1X2* were found in chromosomes of EC0880B_C_, EC0880B_R2_, EC0880B_R3_ and EC0880B_R4_, with 96–98% nucleotide identities and a 100% coverage to the relevant reference sequence. The precise copy number of IS*1X2* could not be determined in the Unicycler short-read-only assembly of EC0880B_R1_, although 100% of this insertion sequence was recovered from five connected nodes (with 96–100% nucleotide identities to the reference sequence) in the assembly graph of this genome. These five nodes had a median read depth of 11 folds, suggesting 11 copies of IS*1X2*, which is consistent with the identified deletion in this isolate’s chromosome (Figure 3c). Similarly, 11 copies of IS*1X2* were estimated from the Unicycler short-read-only assembly graph of EC0880B with ≥95% nucleotide identities.

Interestingly, the other oxygen-insensitive nitroreductase gene *nfsA* in chromosomes of EC0880B_C_, EC0880B_R2_, EC0880B_R3_, and EC0880B_R4_, was interrupted by the same IS*1X2* element that was associated with the deletion of *nfsB*, and locations of these interruptions were the same as that previously observed in EC0880B [7]. The same interruption of *nfsA* was seen in EC0880B_R1_, albeit the sequence of the interruptive IS*1X2* could not be reliably recovered from the short-read-only assembly graph of this genome.

## Discussion

We confirmed monoclonal nitrofurantoin heteroresistance in *E. coli* blood strains EC0026B and EC0880B using PAP, the gold standard for detecting heteroresistance [29]. The MIC of each strain was 8–16 times its maximum non-inhibitory nitrofurantoin concentration. Each strain had a small subpopulation (2.14×10^−6^–9.85×10^−5^) that could grow at 32 mg/L (½ MIC). This frequency of less susceptible cells in each strain was greater than a proposed minimum of 10^−7^ for defining heteroresistance [29]. Since no genetic variation in *nfsA* or *ribE* was identified between isolates of each strain, and neither EC0026B nor EC0880B produces the multidrug-efflux pump OqxAB [7], we attribute the observed nitrofurantoin heteroresistance to the IS*1*-associated deletion of *nfsB* regions. This kind of deletion probably occurred in a UTI-associated *E. coli* ST131 strain, whose nitrofurantoin MIC increased from 8 mg/L to 128 mg/L upon intermittent *in vivo* exposures to a therapeutic dose of nitrofurantoin in six months [30, 31].

We also show that an IS*1* element, IS*1X2*, was associated with the truncation of capsule locus *wzi-wza-wzb-wzc* in EC0880B_C_. All deletion events observed in our study are consistent with the abortive transposition model of IS*1*-mediated deletion, where a DNA duplex nicks at an end of IS*1* and ligates to a target site on the same molecule, causing a loss of DNA between these two breakpoints [32]. Moreover, IS*1X2* had interrupted *nfsA* in parental isolate EC0880B, and we have previously elucidated that IS*1R* interrupted *nfsA* in two nitrofurantoin-resistant isolates [7]. Taken together, IS*1*-associated deletion and interruption of *nfsA* or *nfsB* reduces nitrofurantoin susceptibility of *E. coli* and generally, may occur at other loci, resulting in significant genomic and possibly phenotypic changes. As such, we emphasise the importance of monitoring the prevalence and genomic locations of IS*1* using long-read sequencing.

Nitrofurantoin heteroresistance in *E. coli* strain K-12 MG1655 was reported earlier, with a small subpopulation growing at a maximum nitrofurantoin concentration of 8 mg/L while the majority of cells had an MIC of 4 mg/L [33]. Our PAP experiment discovered similar heteroresistance in *E. coli* strain ATCC 25922, which showed a maximum non-inhibitory nitrofurantoin concentration of 4 mg/L and a subpopulation growing at 8 mg/L with an average frequency of 9% (Figure 2a). Nonetheless, neither the level of the reduction in nitrofurantoin susceptibility nor the nitrofurantoin MIC of each strain is as great as those of strains EC0026B and EC0880B. Because both EC0026B and EC0880B had an MIC of 64 mg/L, which is the EUCAST clinical breakpoint for nitrofurantoin resistance (>64 mg/L), these strains may grow under therapeutic or prophylactic dosing of nitrofurantoin [34, 35] and gain adaptive mutations under sub-MIC exposure [36] in urinary tract or gut. Moreover, the deletion of *nfsB* in these strains would cause an irreversible reduction in their nitrofurantoin susceptibility, paving the way to the emergence of nitrofurantoin-resistant cells if there were to be subsequent resistance mutations, such as those in *nfsA, ribE*, and the *marA*-*marB* intergenic region [36, 37]. Importantly however, because the loss of *nfsB* reduces the reproduction rate of *E. coli* [30], the establishment of *ΔnfsB* mutants may only occur under selective pressure of nitrofurantoin.

We consider IS*1*-associated nitrofurantoin heteroresistance as a potential threat to the management of UTI for three reasons. First, the frequency of the subpopulation in strain EC0026B and EC0880B growing at 0.5× MIC (2–99 cells per million) was 10–100 times reported frequencies of spontaneous resistance-conferring point mutations in *E. coli* [38, 39]. An early study shows a 30–2000 folds increase in the deletion frequency when IS*1* is present [40]. Second, IS*1* provides *E. coli* with adaptive fitness. Specifically, in the absence of nitrofurantoin, the nitrofurantoin-susceptible major population (the wild type) can transmit and establish colonisation or infection as homogeneously susceptible *E. coli* does and likely outcompete nitrofurantoin-resistant *E. coli* that has lost *nfsA* and/or *nfsB*; and in the presence of nitrofurantoin, the infection/colonisation may persist with the subpopulation of reduced susceptibility. Third, nitrofurantoin-heteroresistant isolates may not be detected by routine susceptibility tests in clinical settings if the less-susceptible subpopulation has a low frequency. To determine the mechanism of nitrofurantoin heteroresistance in *E. coli*, our study took advantage of hybrid *de novo* assemblies in accurately resolving complete bacterial genomes and identifying structural variation. Further work is needed to elucidate how IS*1* is associated with genetic deletions and to compare fitness between mutants and parental isolates. Heightened surveillance for prevalence and insertion sites of IS*1* in clinical *E. coli* isolates will help us to understand how insertion sequences contribute to bacterial evolution.

## Conclusions

*E. coli* can be stably heteroresistant to nitrofurantoin with a potential clinical significance, and such heteroresistance may be a precursor of nitrofurantoin resistance. IS*1*-family elements are associated with large-scale deletion of genomic regions containing *nfsB* and are potentially important agents of genomic and phenotypic changes in *E. coli*.

## Acknowledgement

We thank Dr Bruno Pichon at the UK Health Security Agency (formerly, Public Health England) for facilitating WGS of *E. coli* isolates. YW, EJ, CSB, MJE, and SS acknowledge the National Institute for Health Research Health Protection Research Unit in Healthcare Associated Infections and Antimicrobial Resistance at Imperial College London in partnership with the UK Health Security Agency, in collaboration with, Imperial Healthcare Partners, University of Cambridge and University of Warwick [grant number NIHR200876]. YW is an Imperial Institutional Strategic Support Fund Springboard Research Fellow, funded by the Wellcome Trust and Imperial College London [grant number PSN109]. EJ is an Imperial College Research Fellow, funded by the Rosetrees Trust and Stoneygate Trust [grant number M683].

## Transparency declarations

None to declare.

## References

1. National Institute for Health and Care Excellence. Guidance, NICE advice and quality standards (NG109, NG112, NG113). NICE. https://www.nice.org.uk/guidance/published (2018, accessed 13 March 2023).

2. Mottaghizadeh F, Mohajjel Shoja H, Haeili M, Darban-Sarokhalil D. Molecular epidemiology and nitrofurantoin resistance determinants from nitrofurantoin non-susceptible Escherichia coli isolated from urinary tract infections. Journal of Global Antimicrobial Resistance 2020;21:335–339.

3. Ho P-L, Ng K-Y, Lo W-U, Law PY, Lai EL-Y, et al. Plasmid-Mediated OqxAB Is an Important Mechanism for Nitrofurantoin Resistance in Escherichia coli. Antimicrobial agents and chemotherapy 2015;60:537–543.

4. Abernethy J, Guy R, Sheridan EA, Hopkins S, Kiernan M, et al. Epidemiology of Escherichia coli bacteraemia in England: results of an enhanced sentinel surveillance programme. Journal of Hospital Infection 2017;95:365–375.

5. Ny S, Edquist P, Dumpis U, Gröndahl-Yli-Hannuksela K, Hermes J, et al. Antimicrobial resistance of Escherichia coli isolates from outpatient urinary tract infections in women in six European countries including Russia. Journal of Global Antimicrobial Resistance 2019;17:25–34.

6. UK Health Security Agency. English surveillance programme for antimicrobial utilisation and resistance (ESPAUR) Report 2021 to 2022 Annexe. https://assets.publishing.service.gov.uk/government/uploads/system/uploads/attachment_d ata/file/1118730/ESPAUR-report-2021-2022-annexe.pdf (2022, accessed 13 March 2023).

7. Wan Y, Mills E, Leung RCY, Vieira A, Zhi X, et al. Alterations in chromosomal genes nfsA, nfsB, and ribE are associated with nitrofurantoin resistance in Escherichia coli from the United Kingdom. Microbial Genomics 2021;7:000702.

8. Jauneikaite E, Honeyford K, Blandy O, Mosavie M, Pearson M, et al. Bacterial genotypic and patient risk factors for adverse outcomes in Escherichia coli bloodstream infections: a prospective molecular epidemiological study. Journal of Antimicrobial Chemotherapy 2022;77:1753–1761.

9. European Committee on Antimicrobial Susceptibility Testing. EUCAST Quality Control. EUCAST. https://www.eucast.org/ast_of_bacteria/quality_control (2022, accessed 27 December 2022).

10. EUCAST. Clinical breakpoints and dosing of antibiotics (v13.0). https://www.eucast.org/clinical_breakpoints.

11. Bolger AM, Lohse M, Usadel B. Trimmomatic: a flexible trimmer for Illumina sequence data. Bioinformatics 2014;30:2114–2120.

12. De Coster W, D’Hert S, Schultz DT, Cruts M, Van Broeckhoven C. NanoPack: visualizing and processing long-read sequencing data. Bioinformatics 2018;34:2666–2669.

13. Vaser R, Šikić M. Time- and memory-efficient genome assembly with Raven. Nat Comput Sci 2021;1:332–336.

14. Wick RR, Judd LM, Gorrie CL, Holt KE. Unicycler: Resolving bacterial genome assemblies from short and long sequencing reads. PLOS Computational Biology 2017;13:e1005595.

15. Wick RR, Holt KE. Polypolish: Short-read polishing of long-read bacterial genome assemblies. PLOS Computational Biology 2022;18:e1009802.

16. Zimin AV, Marçais G, Puiu D, Roberts M, Salzberg SL, et al. The MaSuRCA genome assembler. Bioinformatics 2013;29:2669–2677.

17. Wick RR, Judd LM, Holt KE. Assembling the perfect bacterial genome using Oxford Nanopore and Illumina sequencing. PLOS Computational Biology 2023;19:e1010905.

18. Seemann T. Prokka: rapid prokaryotic genome annotation. Bioinformatics 2014;30:2068–2069.

19. Siguier P, Perochon J, Lestrade L, Mahillon J, Chandler M. ISfinder: the reference centre for bacterial insertion sequences. Nucleic Acids Research 2006;34:D32–D36.

20. Li H. Minimap2: pairwise alignment for nucleotide sequences. Bioinformatics 2018;34:3094–3100.

21. Danecek P, Bonfield JK, Liddle J, Marshall J, Ohan V, et al. Twelve years of SAMtools and BCFtools. GigaScience 2021;10:giab008.

22. Barrick JE, Colburn G, Deatherage DE, Traverse CC, Strand MD, et al. Identifying structural variation in haploid microbial genomes from short-read resequencing data using breseq. BMC Genomics 2014;15:1039.

23. Marçais G, Delcher AL, Phillippy AM, Coston R, Salzberg SL, et al. MUMmer4: A fast and versatile genome alignment system. PLOS Computational Biology 2018;14:e1005944.

24. Cingolani P, Platts A, Wang LL, Coon M, Nguyen T, et al. A program for annotating and predicting the effects of single nucleotide polymorphisms, SnpEff. Fly 2012;6:80–92.

25. Carver T, Harris SR, Berriman M, Parkhill J, McQuillan JA. Artemis: an integrated platform for visualization and analysis of high-throughput sequence-based experimental data. Bioinformatics 2011;28:464–469.

26. Alikhan N-F, Petty NK, Ben Zakour NL, Beatson SA. BLAST Ring Image Generator (BRIG): simple prokaryote genome comparisons. BMC Genomics 2011;12:402.

27. Siguier P, Gourbeyre E, Varani A, Ton-Hoang B, Chandler M. Everyman’s Guide to Bacterial Insertion Sequences. Microbiology Spectrum 2015;3:3.2.25.

28. Rahn A, Beis K, Naismith JH, Whitfield C. A Novel Outer Membrane Protein, Wzi, Is Involved in Surface Assembly of the Escherichia coli K30 Group 1 Capsule. Journal of Bacteriology 2003;185:5882–5890.

29. Andersson DI, Nicoloff H, Hjort K. Mechanisms and clinical relevance of bacterial heteroresistance. Nature Reviews Microbiology 2019;17:479–496.

30. Vallée M, Harding C, Hall J, Aldridge PD, Tan A. Exploring the in situ evolution of nitrofurantoin resistance in clinically derived uropathogenic Escherichia coli isolates. Journal of Antimicrobial Chemotherapy 2022;dkac398.

31. Mowbray C, Tan A, Vallée M, Fisher H, Chadwick T, et al. Multidrug-resistant Uro-associated Escherichia coli Populations and Recurrent Urinary Tract Infections in Patients Performing Clean Intermittent Self-catheterisation. European Urology Open Science 2022;37:90–98.

32. Grindley NDF, Sherratt DJ. Sequence Analysis at IS1 Insertion Sites: Models for Transposition. Cold Spring Harb Symp Quant Biol 1979;43:1257–1261.

33. Roemhild R, Linkevicius M, Andersson DI. Molecular mechanisms of collateral sensitivity to the antibiotic nitrofurantoin. PLOS Biology 2020;18:e3000612.

34. Muller AE, Verhaegh EM, Harbarth S, Mouton JW, Huttner A. Nitrofurantoin’s efficacy and safety as prophylaxis for urinary tract infections: a systematic review of the literature and meta-analysis of controlled trials. Clinical Microbiology and Infection 2017;23:355–362.

35. Huttner A, Wijma RA, Stewardson AJ, Olearo F, Von Dach E, et al. The pharmacokinetics of nitrofurantoin in healthy female volunteers: a randomized crossover study. Journal of Antimicrobial Chemotherapy 2019;74:1656–1661.

36. Pereira C, Warsi OM, Andersson DI. Pervasive Selection for Clinically Relevant Resistance and Media Adaptive Mutations at Very Low Antibiotic Concentrations. Molecular Biology and Evolution 2023;40:msad010.

37. Sandegren L, Lindqvist A, Kahlmeter G, Andersson DI. Nitrofurantoin resistance mechanism and fitness cost in Escherichia coli. Journal of Antimicrobial Chemotherapy 2008;62:495–503.

38. Kram KE, Finkel SE. Culture Volume and Vessel Affect Long-Term Survival, Mutation Frequency, and Oxidative Stress of Escherichia coli. Applied and Environmental Microbiology 2014;80:1732–1738.

39. Liu X, E. Painter R, Enesa K, Holmes D, Whyte G, et al. High-throughput screening of antibiotic-resistant bacteria in picodroplets. Lab on a Chip 2016;16:1636–1643.

40. Reif HJ, Saedler H. IS1 is involved in deletion formation in the gal region of E. coli K12. Molec gen Genet 1975;137:17–28.

