## Supplementary tables for "IS*1*-related large-scale deletion of chromosomal regions harbouring oxygen-insensitive nitroreductase gene *nfsB* causes nitrofurantoin heteroresistance in *Escherichia coli*"

April 2023

### Supplementary tables

**Supplementary Table 1.** Accession numbers of *E. coli* whole-genome sequencing data stored under BioProject PRJEB58678 in the European Nucleotide Archive.

| Strain | Isolate | Sample | Illumina reads | MinION reads | Assembly |
|---|---|---|---|---|---|
| EC0026B | EC0026B$_{C}$ | ERS14863249 | ERR11181256 | ERR11181589 | GCA_949786325 |
| | EC0026B$_{R1}$ | ERS14863250 | ERR11181257 | ERR11181590 | GCA_949786345 |
| | EC0026B$_{R2}$ | ERS14863251 | ERR11181258 | ERR11181591 | GCA_949786305 |
| | EC0026B$_{R3}$ | ERS14863252 | ERR11181259 | ERR11181592 | GCA_949786315 |
| | EC0026B$_{R4}$ | ERS14863253 | ERR11181260 | ERR11181593 | GCA_949786285 |
| EC0880B | EC0880B$_{C}$ | ERS14863254 | ERR11181261 | ERR11181594 | GCA_949786355 |
| | EC0880B$_{R1}$ | ERS14863255 | ERR11181262 | Not applicable | GCA_949786335 |
| | EC0880B$_{R2}$ | ERS14863256 | ERR11181263 | ERR11181595 | GCA_949786365 |
| | EC0880B$_{R3}$ | ERS14863257 | ERR11181264 | ERR11181596 | GCA_949786295 |
| | EC0880B$_{R4}$ | ERS14863258 | ERR11181265 | ERR11181597 | GCA_949786275 |

**Supplementary Table 2.** Summary of *E. coli* isolates and complete sequences (all circular) recovered from *de novo* genome assemblies. Plus signs indicate isolates for which MinION reads were available, and asterisks indicate chromosome sequences used as references for comparison. No complete chromosome or plasmid sequences were recovered for EC0880B$_{R1}$ because DNA extracted from this isolate was inadequate for MinION sequencing.

| Strain | Isolate | MinION | Chromosome length | Plasmid length |
|---|---|---|---|---|
| EC0026B | EC0026B$_{C}$ | + | 4,741,476 bp * | 88,489 bp; 73,423 bp; 1,788 bp |
| | EC0026B$_{R1}$ | + | 4,730,372 bp | 88,654 bp; 73,432 bp; 1,787 bp |
| | EC0026B$_{R2}$ | + | 4,721,705 bp | 88,510 bp; 73,423 bp |
| | EC0026B$_{R3}$ | + | 4,722,421 bp | 88,442 bp; 73,423 bp |
| | EC0026B$_{R4}$ | + | 4,725,107 bp | 88,429 bp; 73,423 bp |
| EC0880B | EC0880B$_{C}$ | + | 4,837,144 bp * | 159,448 bp; 33,743 bp |
| | EC0880B$_{R1}$ | | Not recovered | 33,743 bp; 1,783 bp |
| | EC0880B$_{R2}$ | + | 4,817,091 bp | 159,448 bp; 33,743 bp |
| | EC0880B$_{R3}$ | + | 4,817,712 bp | 159,448 bp; 33,743 bp; 1,783 bp |
| | EC0880B$_{R4}$ | + | 4,800,993 bp | 159,448 bp; 33,743 bp |
